# Tumor heterogeneity assessed by sequencing and fluorescence *in situ* hybridization (FISH) data

**DOI:** 10.1101/2020.02.29.970392

**Authors:** Haoyun Lei, E. Michael Gertz, Alejandro A. Schäffer, Xuecong Fu, Yifeng Tao, Kerstin Heselmeyer-Haddad, Irianna Torres, Xulian Shi, Kui Wu, Guibo Li, Liqin Xu, Yong Hou, Michael Dean, Thomas Ried, Russell Schwartz

## Abstract

Computational reconstruction of clonal evolution in cancers has become a crucial tool for understanding how tumors initiate and progress and how this process varies across patients. The field still struggles, however, with special challenges of applying phylogenetic methods to cancers, such as the prevalence and importance of copy number alteration (CNA) and structural variation (SV) events in tumor evolution, which are difficult to profile accurately by prevailing sequencing methods in such a way that subsequent reconstruction by phylogenetic inference algorithms is accurate. In the present work, we develop computational methods to combine sequencing with multiplex interphase fluorescence in situ hybridization (miFISH) to exploit the complementary advantages of each technology in inferring accurate models of clonal CNA evolution accounting for both focal changes and aneuploidy at whole-genome scales. We demonstrate on simulated data that incorporation of FISH data substantially improves accurate inference of focal CNA and ploidy changes in clonal evolution from deconvolving bulk sequence data. Analysis of real glioblastoma data for which FISH, bulk sequence, and single cell sequence are all available confirms the power of FISH to enhance accurate reconstruction of clonal copy number evolution in conjunction with bulk and optionally single-cell sequence data.

**Availability:** github.com/CMUSchwartzLab/FISH_deconvolution

## 1 Introduction

Cancer progression has long been understood to be driven by clonal evolution [25], but our understanding of the mechanisms and implications of that observation are currently undergoing a dramatic revision. This growing insight has been driven largely by two key innovations: the advance of high-throughput sequencing methods to characterize tumor genomics with ever-finer precision and accuracy [23] and the concurrent advance of computational biology methods that have made it possible to interpret those sequencing data to tell coherent stories about how individual cancers or the space of all cancers collectively develop [3]. A crucial component of those latter advances has been the development of the field of tumor phylogenetics [28], which develops computational methods to construct models of evolution in cancers from tumor genomic data.

Methods for clonal phylogenetics have attracted great interest in computational biology circles, especially as the field has come to a greater understanding of the complexity of tumor evolution mechanisms and the algorithmic challenges of reconstructing them from available genomic data. A particular area of recent interest in this regard has been development of better methods for resolving evolution by copy number alterations (CNAs) and the structural variations (SVs) that produce them. While the importance of CNAs and SVs in cancer has long been known [37] and some of the first methods for clonal lineage reconstruction focused on CNA-driven evolution [27], much of the tumor phylogeny field has focused historically on single nucleotide variants (SNVs), with CNAs omitted (e.g., [34]) or treated largely as a confounding factor for inferring SNV-driven evolution (e.g.,[16]). Relatively few computational methods have been created to date for the purpose of inferring tumor evolution by CNAs, either singly [33,29,17,15] or jointly with SNVs [20], and it is only recently that methods have begun to appear for capturing evolution by SVs more broadly [14]. Yet the biological evidence over the same time has strongly indicated that CNVs, and the SVs producing them, outperform SNVs and other focal changes in predicting treatment response [30] and are likely the dominant mechanism by which tumors develop and functionally adapt to escape controls on cell growth [37].

CNA-driven evolution creates substantial complications relative to SNV-driven evolution. In part, modeling CNAs is a challenge because it is a less commonly studied kind of mechanism in phylogenetics in general and thus requires algorithmic innovations. In part, the challenge is inherent to the problem. CNAs create particular complications because they can occur on multiple scales with sometimes overlapping variations that can be difficult to resolve. CNAs are also particularly challenging for deconvolutional approaches to phylogenetics [2] — which involve computationally separating mixtures of clones from bulk sequence data and which remain a necessity for the field due to the scarsity of large cohorts with single-cell data — because the basic deconvolution problem on CNAs is underdetermined without additional data or problem constraints. CNA methods also have particular difficulty dealing with ploidy changes, particularly via whole-genome duplication (WGD), because ploidy is difficult to infer accurately from sequence data alone. While recent methods have shown it to be possible to perform accurate CNA construction using multi-region bulk sequencing [35] or single-cell sequencing [36], these methods require limiting assumptions, e.g., that WGD can occur only once in a tumor’s history. Furthermore, large cohorts with multi-region bulk or single-cell sequencing are still lacking and it remains an open question how best to perform large-scale tumor genomic studies that will be informative for clonal CNA evolution.

The problem of accurately reconstructing the process of ploidy change in tumor evolution is concerning, partly because WGD is now recognized as a statistical marker of aggressive cancers [26,4] but this observation lacks a clear biological mechanism. Earlier models of WGD in tumor evolution, which proposed a single early WGD event as a prerequisite for tumorigenesis in chromosomally instable cancers [13], are now known to be as excessively simplistic, as WGD is neither necessary nor necessarily a one-time event in a cancer’s evolution [26]. Rather, WGD can be seen to be one of many mutation types active to different degrees in different cancers, shaping the likely trajectory and patient-specific risk of diverse progression processes [6].

The present work develops methods for improving our ability to resolve CNA-driven evolution in cancers, following a strategy of multi-omic data integration. Malicik et al. demonstrated the power of integrating bulk and single-cell sequencing data [22] for improving SNV evolution models, a strategy we previously demonstrated successful for CNA-driven evolution as well [21]. Here, we explore the potential of bringing in an additional form of data, multiplex interphase fluorescence in situ hybridization (miFISH), which provides a way to profile tumor evolution in single cells at small numbers of probes [18] without normalization artifacts that make ploidy a challenge for purely sequence-based studies. While miFISH limits one to just a few copy number markers per cell, its easy scalability to large numbers of cells has made it powerful tool for CNV tumor phylogenetics in its own right especially when the FISH probes are placed strategically at loci that are recurrently amplified in the tumor type of interest [27,10,8,9,38].

Here, we develop a new method for integrating bulk sequence with singlecell sequence (SCS) and/or miFISH in order to combine advantages of each technology for improved reconstruction of copy number evolution at the single cell level. We show with semi-simulated data that these two kinds of data each contribute in distinct and synergistic ways to more accurate inference of CNA-driven evolution, especially in aneuploid tumors, and demonstrate their practical value on a study of glioblastoma profiled by bulk, SCS, and miFISH. Together, the work demonstrates the value of bringing miFISH or related methods for cytometric analysis into sequence-based tumor phylogeny studies if we are to accurately reconstruct mechanisms of CNA-driven evolution in cancers.

## 2 Methods

### 2.1 Objective function

While the mixed membership model in our previous work [21] is still suitable to describe our problem, the previous objective function is too simple for our new problem, since it does not include FISH data. The FISH data is relevant only at a few loci, but with an appropriate objective function allows us to estimate the genome-wide ploidy and thereby to inform the analysis of unnormalized copy number at all loci. We therefore designed a new objective function for the problem intended to capture information from bulk copy numbers, miFISH copy numbers, and single cell sequencing (SCS) data, as well as phylogenetic constraints:

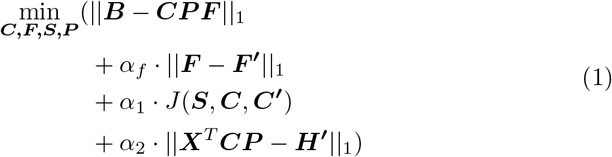

where the desired outputs are a matrix of inferred normalized copy numbers ***C*** of probes across the genome, a diagonal matrix ***P*** with the inferred ploidy divided by *2* for each in the diagonal, and an inferred clonal frequency matrix ***F. CP***, then transforms the *normalized cells* with mean copy number two to *unnormalized cells* with putative absolute copy numbers. As explained below, we require ***C*** to be integral, but not ***C′***.

The potential inputs are **F*′***, a matrix of reference mixture fraction information derived from miFISH data; ***C′***, an optional matrix of normalized copy numbers of input SCS data; ***H′***, the reference copy numbers from miFISH data for the genomic regions covered by FISH probes; and ***X***, a 0-1 matrix identifying copy number segments of the genome covered by each FISH probe. ***S*** is a phylogeny inferred in the process of computing the objective function. ||***B – CPF***||_1_ is the deviation between true and inferred mixed copy number in the bulk tumor. ||***F–F′***||_1_ describes the deviation between inferred fraction ***F*** and reference fraction ***F′, J***(***S, C, C′***) is the cost of the phylogeny that is built on inferred cell clones ***C*** and reference cell clones ***C′. X*** is a sparse matrix such that ***X**_ij_* = 1 if *i* is the index FISH probe in the genomic position of single-cell data and *i* = *Index*[*j*], where each element in *Index* is the index of one FISH probe on the genomic axis of the SCS data. The product ***X***^*T*^***C*** represents a subset of genomic position that contains only the copy number information for segments covered by FISH probes and zeros elsewhere. Then ||***X**^T^**CP*** — ***H′***||_1_ describes the deviation between inferred and reference copy number at loci where FISH probes are located. *α_f_*, *α*_1_ and *α*_2_ are regularization parameters to allow for the balance between deconvolution quality and quality of fit of the inferences to reference mixture fractions, miFISH copy numbers, SCS copy numbers, and a minimum evolution model.

We elaborate in the following subsections on specific constraints involving the terms of the objective function.

#### 2.2 Estimating F

##### 2.2.1 ||*B – CPF*||_1_ constraints

We define the *L*1 distance of ||***B – CPF***||_1_ as:

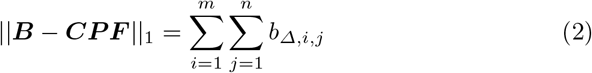

with constraints:

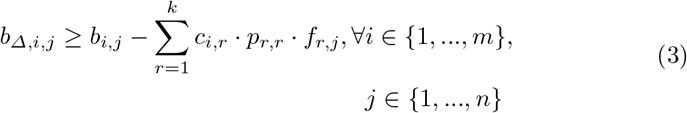

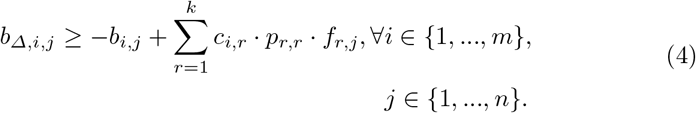

where *m* is the number of total genomic loci, *n* is the number of bulk tumor samples, and *k* is the number of cells.

##### 2.2.2 F constraints

Since ***F*** is a weighted matrix, each column of ***F*** should add up to be 1 and all entries are non-negative. All the entries in ***C*** represent the (average) copy number of a certain interval. As currently implemented, the entries should be non-negative integers, but this is not inherent because the integral copy numbers across the interval may vary leading to a non-integral average copy number.

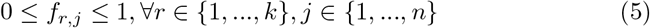

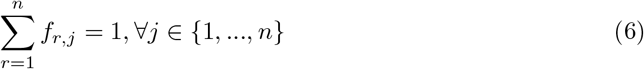

##### 2.2.3 ||*F – F′*||_1_ constraints

We apply *L*1 distance on ||***F – F′***||_1_:

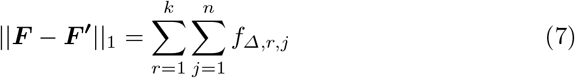

with constraints:

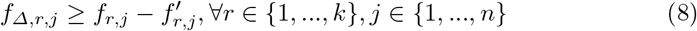

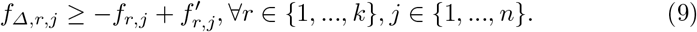

In summary, when we estimate ***F***, we optimize:

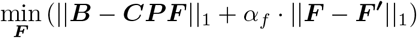

with constraints (3)-(9).

#### 2.3 Estimating S

##### 2.3.1 *J*(*S, C, C′*) constraints

The term *J*(***S, C, C′***) includes a phylogenetic relationship on the inferred cell data ***C*** and the reference cell data ***C′***, we define a phylogenetic structure with a *K* × *K* directed adjacent matrix ***S***, where *K* = 2*k* + 1, the first *k* columns indicate ***C***, the next *k* columns indicate ***C′*** and the last column indicates a root with normalized copy numbers all-2 (diploid). We introduce a vertex set *T* = {1,…, 2*k* + 1} that represents the set of all cells in ***S***. Let *r* be the unique, predetermined, root of *T*. For *t,u,v* ∈ *T*, we introduced the binary variables 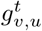 representing the amount of flow along edge (*u, v*) with destination *t* ∈ *T*. Then the full constraints are: flow conservation on the Steiner vertices:

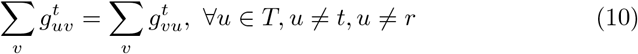

inflow/outflow constraints on terminals in *T*:

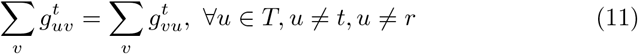

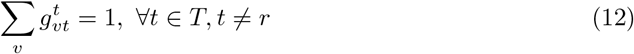

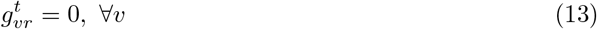

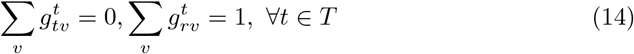

positive flow on an edge iff the edge is selected:

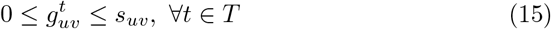

no self loop:

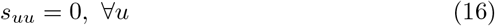

binary variable for 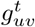 and *s_u,v_*:

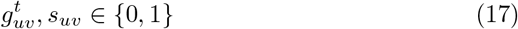

##### 2.3.2 Phylogenetic cost

We then define the measurement for evolutionary distance across each edge (*u, v*) in the tree as *L*1 distance of the copy number profiles of the edge endpoints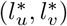 and introduce a minimum evolution model defined by ***S*** to estimate the phylogenetic cost:

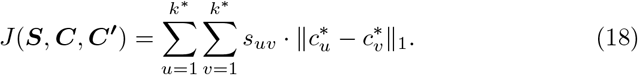

We define the phylogeny objective to be derived from normalized copy numbers, effectively ignoring ploidy changes in the evolution objective and measuring distance from localized focal copy number variations only. One might plausibly improve on this model by accounting for ploidy changes separately as evolutionary events [8,9] or adopting a more nuanced general model of copy number change, such as the MEDICC model [29]. The L1 distance of normalized copy numbers is used as a heuristic because of the difficulty of incorporating these other model types into the ILP framework.

#### 2.4 Estimating C

##### 2.4.1 *C* constraints

We impose some basic constraints on ***C***: (1) all copy numbers are no larger than a certain maximum number *c_max_*, which is set at *10* in our tests; (2) all copy numbers must be integers.

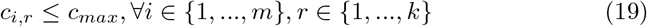

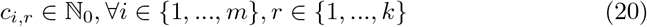

##### 2.4.2 ||*C – C′*||_1_ constraints

The ||***C – C′***||_1_ term is not explicitly expressed in the objective function. Instead, it exists in the *J*(***S, C, C’***) term since we apply *L*_1_ distance between two nodes in S as the edge weight (Eq. 18). Then we redefine:

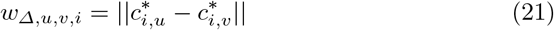

with constraints:

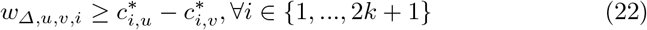

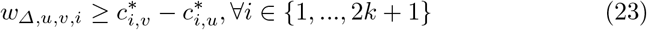

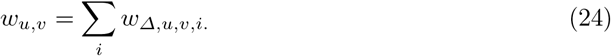

##### 2.4.3 ||*X*^*T*^*CP* – *H′*||_1_ constraints

FISH probes each cover a genomic interval spanned by SCS data, so ||***X***^*T*^***CP*** — ***H′***||_1_ provides a way to favor consistency between FISH and SCS data over these intervals in the coordinate descent optimization. We note that one might optionally weight this objective term to account for varying clonal frequencies of the FISH cells, although we do not do so here. In this step, the known ***H′***, which represents the *unnormalized* FISH probes, provides additional constraints on copy number in ***C***. To get these constraints, first of all, we define an array *Index* describing mapping of FISH probe regions to SCS copy number regions, where each item indicates the index of FISH probe in the single-cell data, and we define ***X*** as follows:

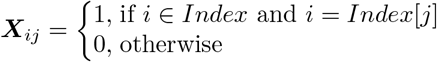

for ∀_*i*_ ∈ {1,…, *m*}, and ∀_*j*_ ∈ {1,…, *Index*.length}

Then we impose *L*1 distance on ||***X**^T^**CP*** – ***H′***|| and redefine it as:

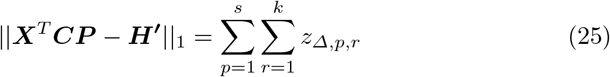

with constraints:

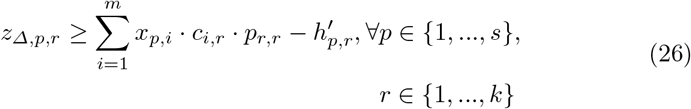

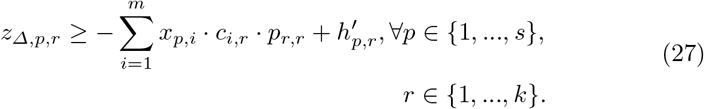

where *s* is the number of FISH probes, *k* is the number of cells.

#### 2.5 Estimating P

##### 2.5.1 *P* constraints

***P*** is the diagonal matrix whose diagonal elements are the half ploidies (rescaling factors) to transform the *normalized copy numbers to* unnormalized copy numbers. We also set lower (*p_min_*) and upper (*p_max_*) bounds for *p_ij_*, and these are set at 0 and 8 respectively in our tests below. The complete constraints are then:

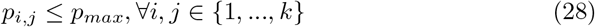

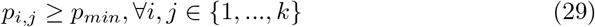

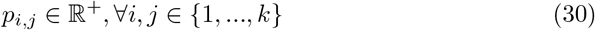

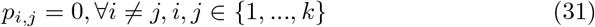

##### 2.5.2 ||*X*^*T*^*CP* – *H′*||_1_ constraints

Unlike in 2.4.3, in this step, ***C*** is known from the computation in previous step to update ***C***, and we would like to update ***P***. *Index* is defined such that ***X^T^*** represents the *normalized* FISH probes, which we can compute after update ***C***, then ***X^T^C*** can be redefined as ***Y***. Then ***YP*** is the *unnormalized* FISH probes. We still impose *L*1 distance between ***YP*** and ***H′*** and redefine it as:

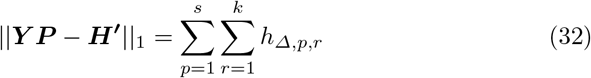

with constraints:

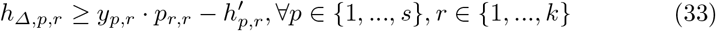

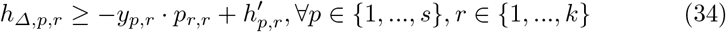

where *s* is the number of FISH probes, *k* is the number of cells.

#### 2.6 Glioblastoma data

We apply the method to SCS and copy number data from two glioblastoma (GBM) patients (GBM07, GBM33) previously described in [21] for which we have samples from 3 tumor regions per patient. In addition, we have FISH data from cells in each region for 8 gene locus probes: *PDGFRA*(4q), *APC*(5q), *EGFR*(7p), *MET*(7q), *MYC*(8q), *CCND1*(11q), *CHEK1*(11q), and *ERG*(21q), several of which were selected because they are sites of recurrent amplifications in GBM [32]. Indeed, both patients have copy numbers over 10 at several loci and patient GBM07 has an extreme amplification with copy numbers possibly over 50, at *PDGFRA.* We continue our practice of setting an upper-bound MAX_C0PY=10 and modifying all observed copy numbers > MAX_COPY to be equal to MAX_C0PY=10 [21]. The methods for designing FISH probes and counting FISH copy numbers have been previously described [18,19,26].

#### 2.7 Simulated data

We further rely on simulated data for validation due to the unavailability of real data with known ground truth. We thus generate synthetic data for which all ground truth properties are known, but with the goal of approximating as well as possible characteristics of the true GBM data described in Sec. 2.6. For this purpose, we generate six data structures per synthetic data set:

1. 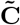: a matrix of normalized copy number profiles of all selected clones, used to compose bulk tumor data
2. 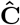: a matrix of normalized copy number profiles of major clones in 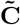, used to evaluate the performance
3. 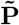: a diagonal matrix of half ploidies of all selected clones
4. 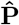: a diagonal matrix of half ploidies of major clones
5. 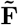: a matrix of mixture fractions of all selected clones in each region
6. 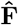: a matrix of mixture fractions of major clones in each region

We use a sample of true single-cell data from the GBM07 dataset as sources of realistic genome-wide normalized copy number vectors, representing SCS data for a set of clones of unknown ploidym then artificially assign ploidies to the data. We perform two versions of this assignment, one with ploidy fixed to diploid for all clones and the other with ploidies sampled randomly to produce 60% diploid clones, 30% tetraploid clones, and equal probability of all other integer ploidies from 1 to 8. We then assign clones to tumor regions, with each region assigned uniformly at random two “major” clones, two “minor” clones, and 23 “tiny” clones, with individual cells sampled from Dirichlet distributions weighted by clone type in the proportions 100:1:0.01. We sample artificial SCS data from these clonal frequencies and optionally add noise in the form of perturbations of copy numbers by ±1 with frequency 10%. We then simulate FISH data from these clones by taking the copy numbers of the two major clones for each region and restricting them to locations where FISH probes appear in the real data. We then extract the six major clones as the reference FISH data and corresponding reference fractions for the optimization. Additional details on the generation of simulated data are omitted here due to space constraints but are provided as supplementary material.

## 3 Results

### 3.1 Evaluation on Simulated Data

#### 3.1.1 No ploidy change

We first evaluate the method on simulated data with no ploidy changes, i.e., all diploid data, to provide a basis for comparison with pure deconvolution and with our previous work [21], which did not explicitly model ploidy. Each test makes use of bulk data and the ||***B – CPF***|| deconvolution objective, but we vary tests by whether or not we use each of the other objective terms — ||***F − F′***||, *J*(***S, C, C′***) and ||***X^T^CP − H′***|| — to determine how they contribute individually or in combination to overall accuracy. As shown in Fig. 1, solving the pure deconvolution problem alone yields poor average accuracy (top right, red bar, Fig. 1), although the results improve substantially when we use true SCS data to initialize the method (top left, blue bar, Fig. 1). This observation is consistent with our prior work [21], although the absolute accuracy of these two variants is worse than in our prior work, perhaps because the current method reduced the maximum number of iterations for the Gurobi solver from 100 to 10.

**Fig. 1.**
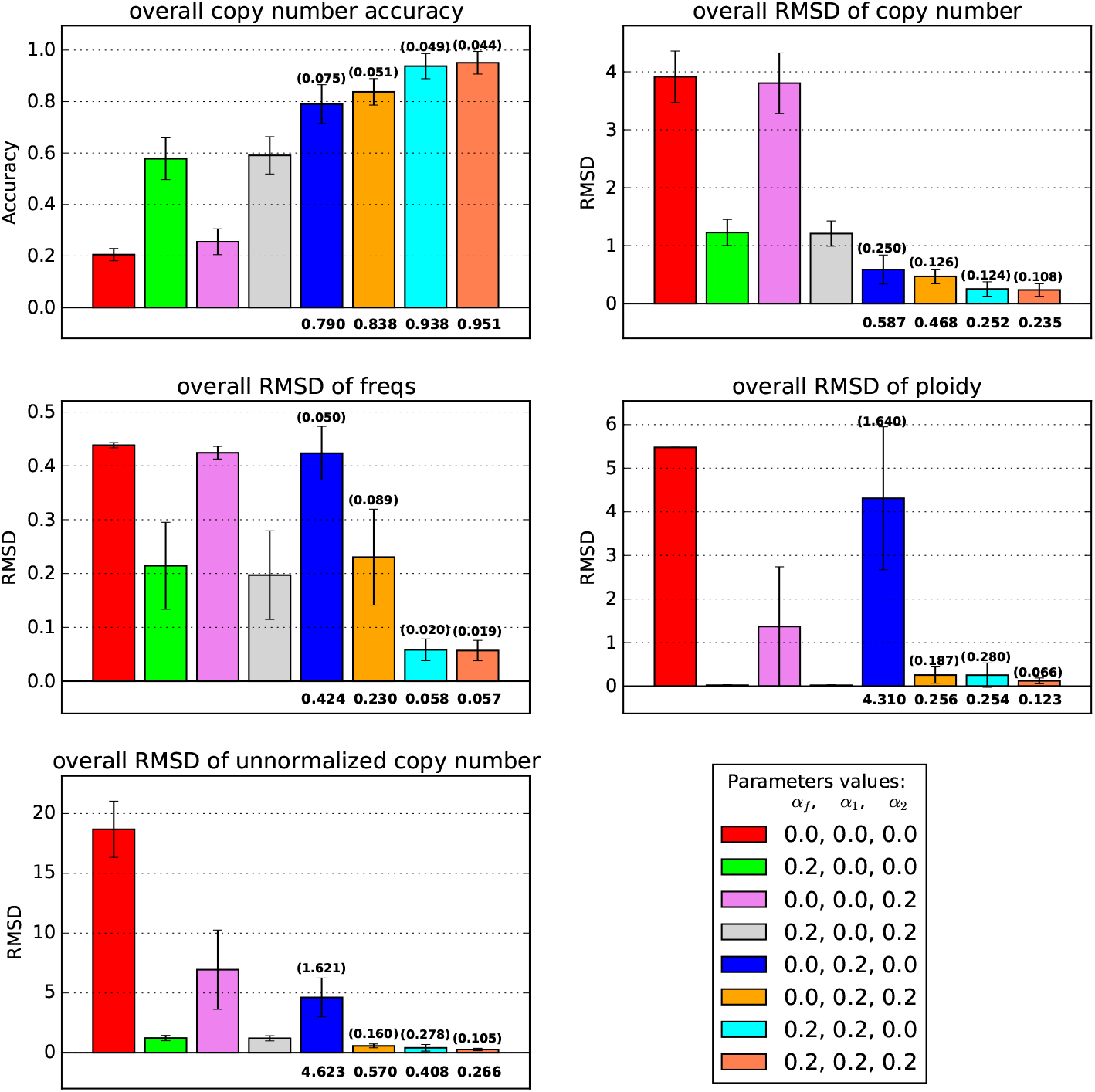
Average accuracy and RMSD of the deconvolution without noise and without ploidy change (n=10). Each bar with different color represents a deconvolution model with different information. In the label table at the bottom right, the numbers represent the value of *α_f_*, *α*_1_, *α*_2_, which are regularization terms for ||***F − F′***||, *J*(***S, C, C′***) and ||***X^T^CP − H′***||, respectively. 0.0 means the corresponding term is not included in the model. The number in parenthesis indicates the standard error of each model, the number under the bar indicate the mean of each model.

Including the term *J*(***S, C, C′***) has a large positive effect on the copy number inference but little impact on the mixture fraction inference (center left, blue bar, Fig. 1). However, adding ||***F – F′***|| to the objective function, and thus using FISH to correct inferred clonal mixture fractions, substantially improves the inference accuracy for both mixture fractions and copy numbers (top left, center left, green bar, Fig. 1). We further found that the combination of the two terms above (*α_f_* = 0.2, *α*_1_ = 0.2, *α*_2_ = 0.0) further improves the performance in both copy number inference and mixture fraction inference (top left, center left, cyan bar, Fig. 1), showing these two components of the objective act in a complementary fashion.

The inclusion of ||***X^T^CP*** — ***H′***|| alone (*α_f_* = 0.0, *α*_1_ = 0.0, *α*_2_ = 0.2), with ||***F − F′***|| (*α_f_* = 0.2, *α*_1_ = 0.0, *α*_2_ = 0.2), or with *J*(***S, C, C′***) (*α_f_* = 0.0, *α*_1_ = 0.2, *α*_2_ = 0.2) yields further improvement (top, violet, grey and orange bars, Fig. 1) in copy number and mixture fraction inference. Including the combination of ||***X^T^CP – H′***|| and *J*(***S, C, C′***) (*α_f_* = 0.0, *α*_1_ = 0.2, *α*_2_ = 0.2) also substantially improves mixture fraction inference (center left, blue and orange bars, Fig. 1) compared to *J*(***S, C, C′***) (*α_f_* = 0.0, *α*_1_ = 0.2, *α*_2_ = 0.0). These results show that the improvements from each objective component and from FISH and SCS data are cumulative. Furthermore, the model with all three penalties (*α_f_* = 0.2, *α*_1_ = 0.2, *α*_2_ = 0.2) yields the best results for mixture fraction inference and both normalized and unnormalized copy number inference (top left and right, center left, and bottom left coral bar, Fig. 1), although it is marginally worse than (*α_f_* = 0.2, *α*_1_ = 0, *α*_2_ = 0.2) for ploidy inference.

We then examine the robustness of the model to noise. As described in Sec. 2.7, we optionally introduced 10% noise to the reference data. The results are similar to those ones without noise and yield qualitatively similar conclusions, although the improvement in the copy number accuracy in the complete model (*α_f_* = 0.2, *α*_1_ = 0.2, *α*_2_ = 0.2) is slightly worse than the model only incorporating FISH and SCS (*α_f_* = 0.2, *α*_1_ = 0, *α*_2_ = 0). Although the model loses some accuracy, it is fairly robust to moderate noise with the current parameters (Fig. 2).

**Fig. 2.**
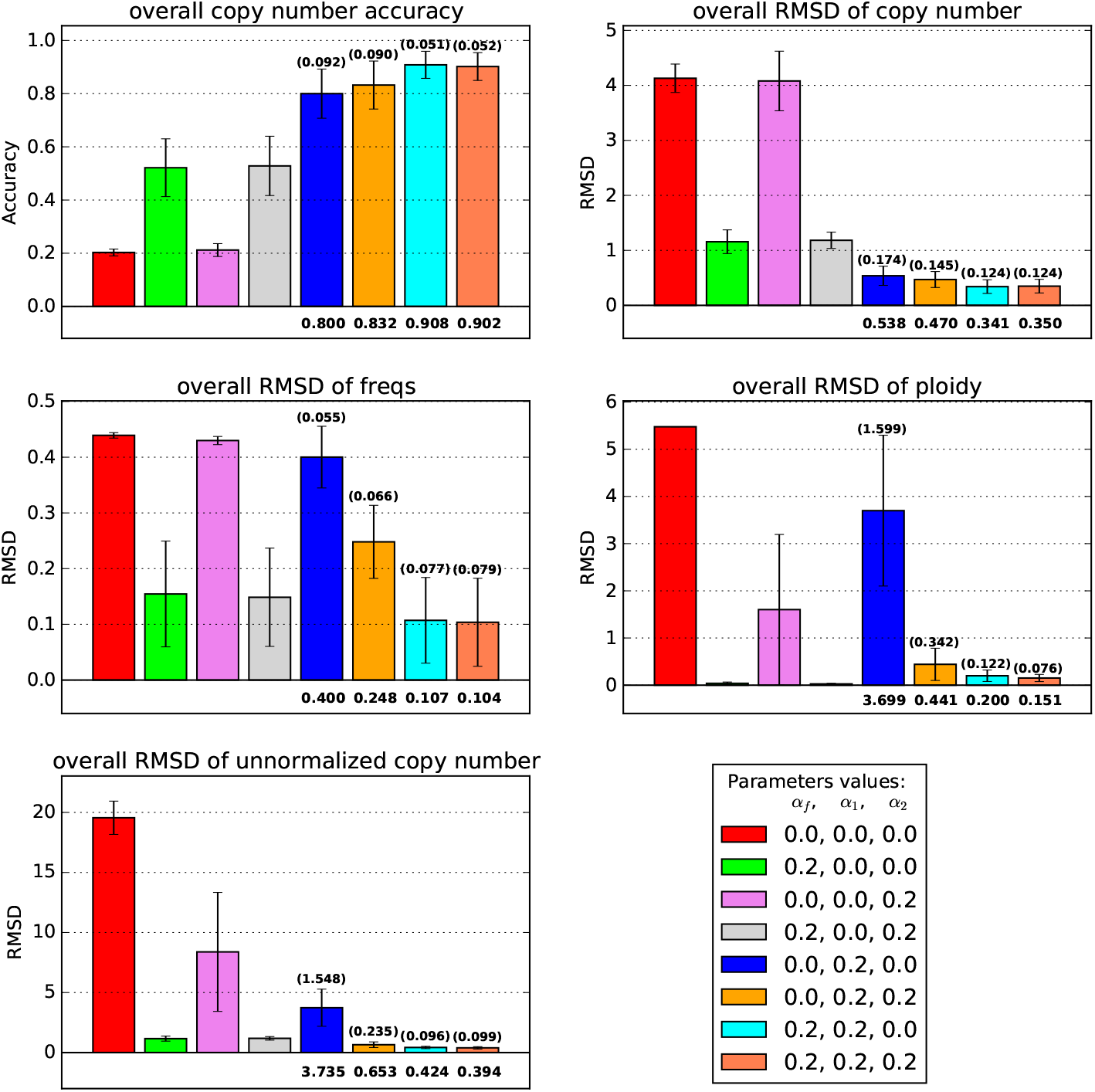
Average accuracy and RMSD of the deconvolution with 10% noise and without ploidy change (n=10). Each bar with different color represents a deconvolution model with different information. In the label table at the bottom right, the numbers represent the value of *α_f_*, *α*_1_, *α*_2_, which are regularization terms for ||***F − F′***||, *J*(***S, C, C′***) and ||***X^T^CP − H′***||, respectively. 0.0 means the corresponding term is not included in the model. The number in parenthesis indicates the standard error of each model, the number under the bar indicate the mean of each model.

#### 3.1.2 With ploidy change

We next examine performance when samples can have variable ploidies. We observe overall lower accuracy of inference across tests when ploidy is variable, although a qualitatively similar profile to the diploid case in Sec. 3.1.1 in how different combinations of objective function terms contribution to accuracy. Pure deconvolution without any information performs much worse without the assumption of diploidy (top left, red bar, Fig. 3). Combining the *J*(***S, C, C′***) and ||***X^T^CP − H′***|| terms, i.e., (*α_f_* = 0.0, *α*_1_ = 0.2, *α*_2_ = 0.2), yields much more obvious improvement when ploidy is variable (top left, blue and orange bar, Fig. 3), though the standard error becomes larger. Furthermore, the model with all three terms yields the best accuracy by all of the measures considered (coral bar, Fig. 3) and in every case, adding in a term of the objective leads to improvement by each measure, showing that each term contributes synergistically to overall accuracy when ploidy is variable.

**Fig. 3.**
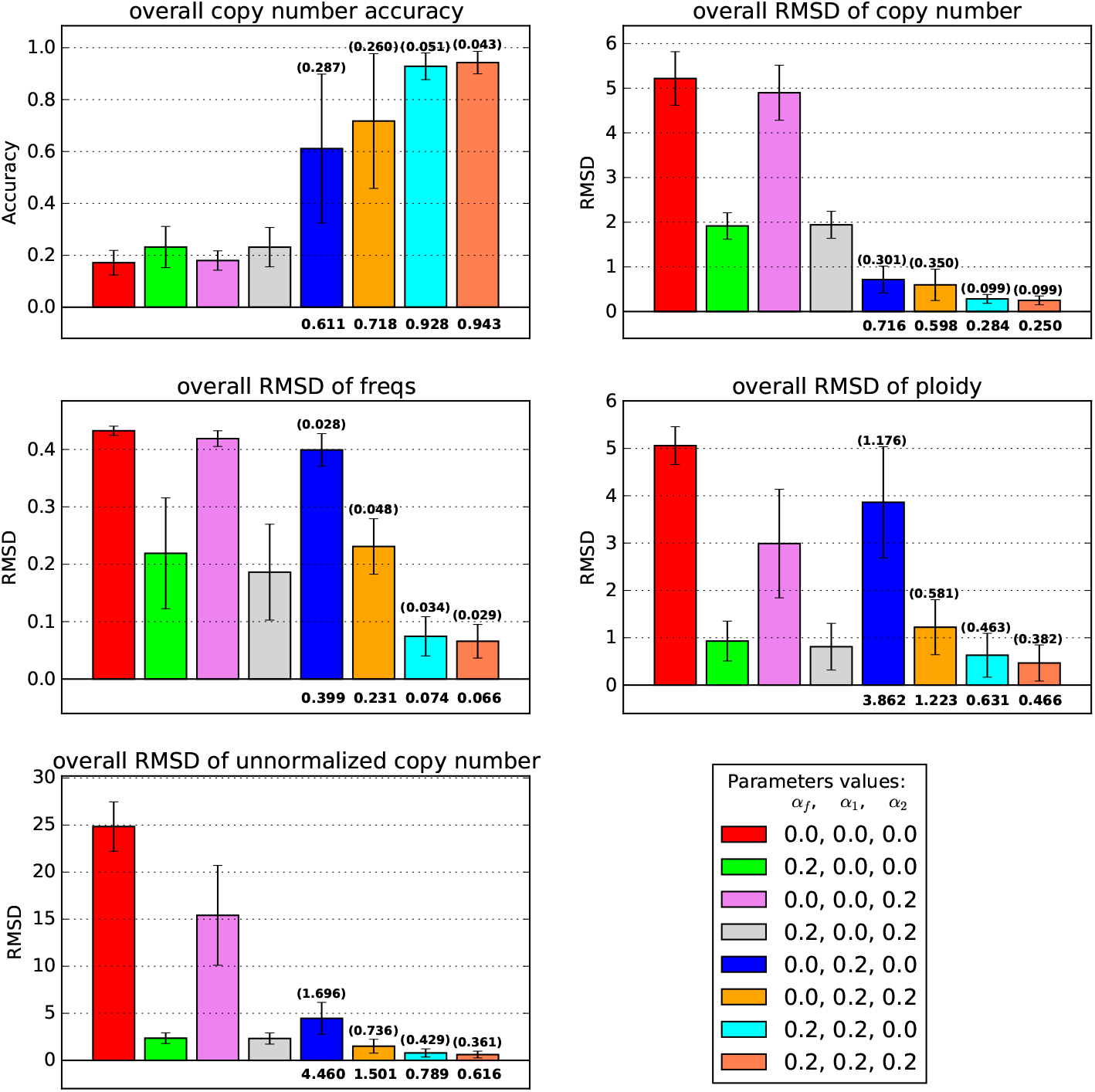
Average accuracy and RMSD of the deconvolution without noise and with ploidy change (n=10). Each bar with different color represents a deconvolution model with different information. In the label table at the bottom right, the numbers represent the value of *α_f_*, *α*_1_, *α*_2_, which are regularization terms for ||***F − F′***||, *J*(***S, C, C′***) and ||***X^T^CP − H′***||, respectively. 0.0 means the corresponding term is not included in the model. The number in parenthesis indicates the standard error of each model, the number under the bar indicate the mean of each model.

When we introduced 10% noise to the data, the conclusion is qualitatively similar (Fig. 4). The complete model (*α_f_* = 0.2, *α*_1_ = 0.2, *α*_2_ = 0.2) again performs the best by all evaluation measurements.

**Fig. 4.**
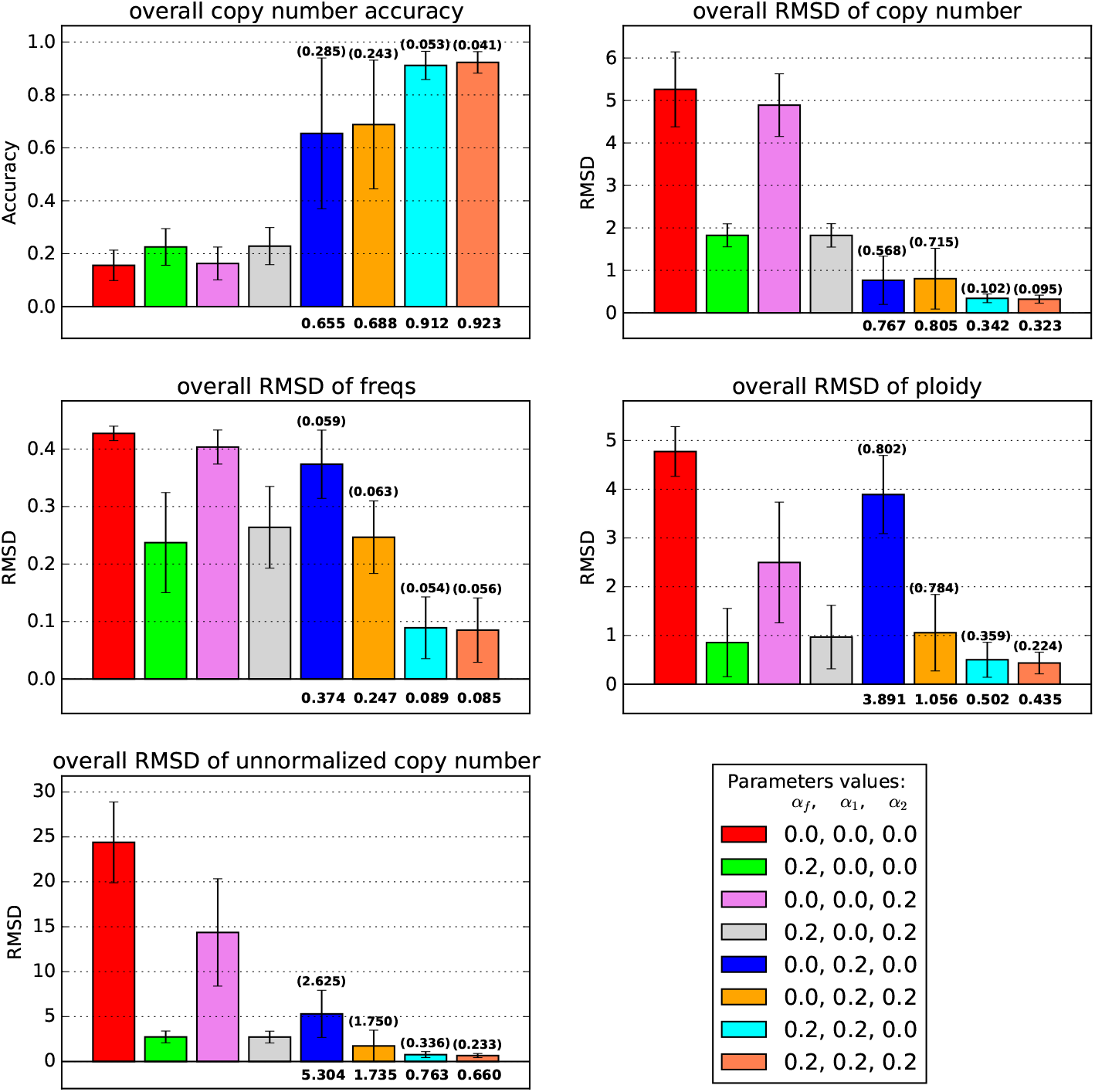
Average accuracy and RMSD of the deconvolution with 10% noise and with ploidy change (n=10). Each bar with different color represents a deconvolution model with different information. In the label table at the bottom right, the numbers represent the value of *α_f_*, *α*_1_, *α*_2_, which are regularization terms for ||***F – F′***||, *J*(***S, C, C′***) and ||***X^T^CP − H′***||, respectively. 0.0 means the corresponding term is not included in the model. The number in parenthesis indicates the standard error of each model, the number under the bar indicate the mean of each model.

#### 3.1.3 Phylogenetic output

Finally, we compare the phylogenetic outputs of the current models. Since the phylogenetic results from the experiments with no ploidy change are trivial (all the ploidies are around 2), we only consider the models with ploidy changes here. We choose the case with the highest overall accuracy of copy number as representatives and plot the phylogenetic trees of three different models that introduce the *J*(***S, C, C′***) term, *J*(***S, C, C′***) term, and ||***X^T^CP − H′***|| term, and all three terms, respectively (Fig. 5). In each case, nodes 0-5 represent the reference cells we observed from the available SCS data, nodes 6-11 represent the inferred cell components constructed by the algorithm, and node 12 represents the assumed diploid root. In each node, we use the notation *NodeIdx;Ploidy* to denote the index of a cell component (cell subclone) and its corresponding ploidy. For example, *12;2* represents the 12^*th*^ cell component (root) and the ploidy of this cell component is 2.

**Fig. 5.**
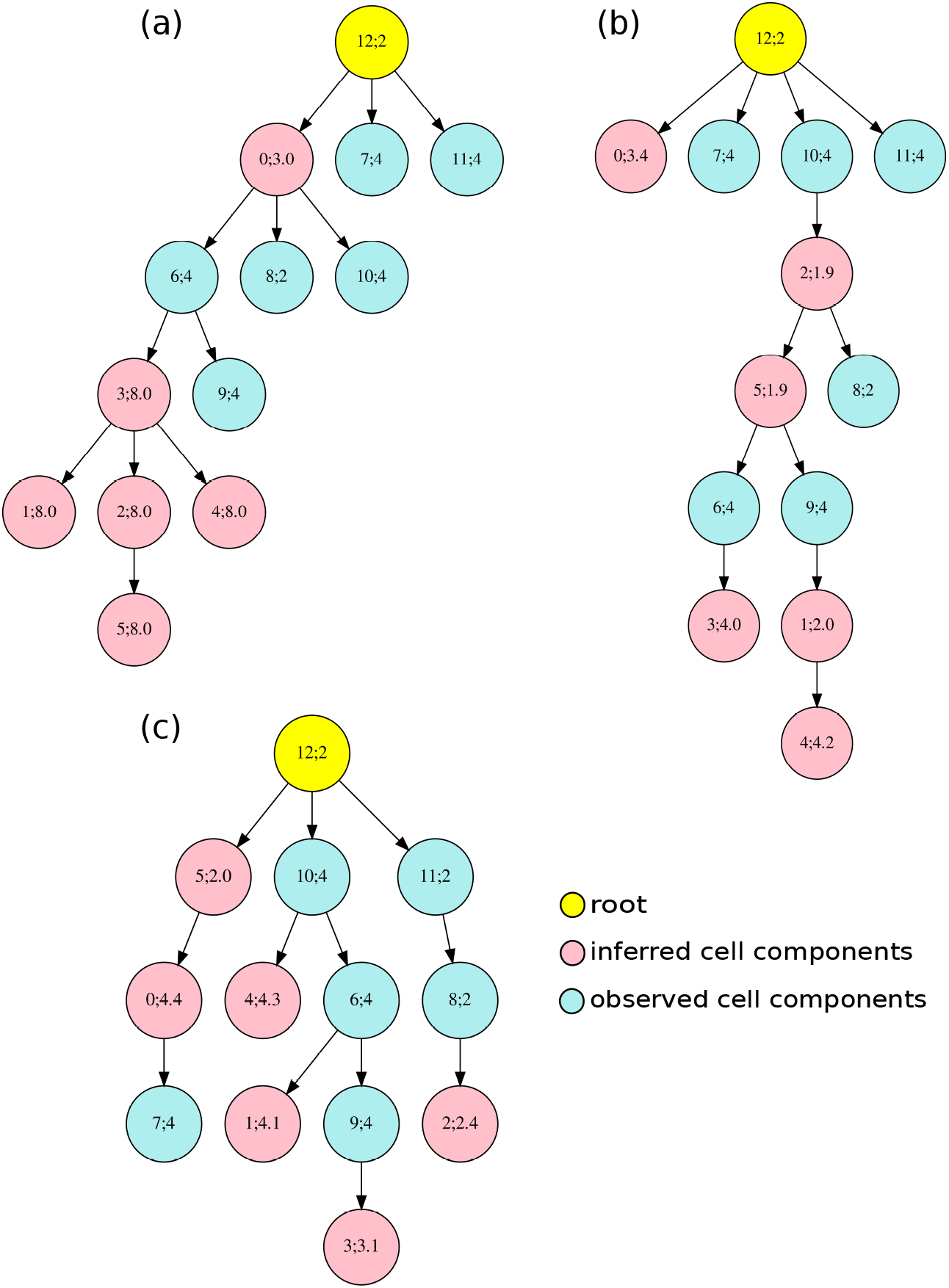
Phylogenetic tree for observed and inferred cell components. The yellow node represents a diploid root cell, the pink nodes are observed cell components and the light blue nodes are inferred cell components. The number pair inside each node provides *Nodeindex; Ploidy*. (a) is the result from the model only including the *J*(***S,C,C′***) term, (b) from the model including *J*(***S,C,C′***) and ||***X^T^CP − H′***||, and (c) from the complete model.

When we only include the *J*(***S, C, C′***) term ((*α_f_* = 0.0, *α*_1_ = 0.2, *α*_2_ = 0.0)), most of the inferred cell components yield unrealistically large copy numbers of 8.0, and the observed and inferred cell components tend to cluster together (Fig. 5 (a)). This may be due to a model tendency to enlarge the ploidy of each inferred cell component to compensate for the deviation between copy number vectors in observed cell components.

When we additionally add the ||***X^T^CP − H′***|| term to update the model ((*α_f_* = 0.0, *α*_1_ = 0.2, *α*_2_ = 0.2)), the ploidy of inferred cell component becomes more realistic, and the inferred and observed cell components show less obvious partitioning (Fig. 5 (b)). We observe that the diploid cell components tend to cluster together (e.g., node 2→node 8) and tetraploid components tend to cluster together (e.g., node 6→node 3). We infer potential WGD events between diploid and tetraploid cell components (e.g. node 5→node 6). This again suggests that the ploidy information from FISH data helps to correct for inferences difficult to make from sequence alone and restore a meaningful phylogenetic structure with ploidy inference among the cell components. The result of introducing ||***F − F′***|| and *J*(***S, C, C′***) together yields a similar pattern (data not shown), suggesting as we might expect that more accurate frequencies can also correct for the ambiguity in inference of *F* and *C* simultaneously that makes the pure copy number deconvolution problem challenging. When we use the complete the model ((*α_f_* = 0.2, *α*_1_ = 0.2, *α*_2_ = 0.2)), the phylogenetic tree becomes more branched, and the diploid and tetraploid cell components are perfectly divided into different branches (node 0→node 7, root→node 10 and root→node 11). Also, the potential WGD events are inferred to happen earlier in the progression (root→node 10 and node 5→node 0). Furthermore, unlike in the previous trees (e.g., node 10→node 2 in Fig. 5 (b)), we see no biologically implausible reversion of WGD events. In addition, although most simulated ploidies in the representative data are tetraploidy, the model is still able to infer a triploidy case (node 9→node 3). All these observations again confirm that the complete model ((*α_f_* = 0.2, *α*_1_ = 0.2, *α*_2_ = 0.2)) not only restores the heterogeneity with best accuracy and performance but also provides the most plausible phylogenetic structure for all the cell components.

### 3.2 Real GBM Data

Finally, we apply the complete model with predefineded parameters ((*α_f_* = 0. 2, *α*_1_ = 0.2, *α*_2_ = 0.2)) on the real glioblastoma cases GBM07 and GBM33. The data include unnormalized copy numbers of bulk sequencing, normalized copy numbers of single-cell sequencing, and unnormalized copy numbers of FISH with estimated ploidies. We apply the *k*-median clustering method on SCS samples and FISH samples to choose *k* = 6 clusters as the reference cells and reference FISH. The copy numbers of bulk samples, profiles of copy number (SCS and FISH) of the cluster centers and profiles of ploidy (FISH) of the cluster centers are the inputs to our the model. We run the method multiple times with different randomly selected real reference data.

Fig. 6 shows typical representative solutions for each case. Since in the real SCS samples, we do not have the true ploidy information, we use “?” to label the ploidy in observed cell components ((c) and (f) in Fig. 6). We first focus on GBM07 case (Fig. 6, (a)-(c)). We observe a pattern of focal CNAs consistent with those described previously in [21] based only on bulk and SCS data. Previous work showed that glioblastomas tend to display at least some chromosome-scale CNAs, such as chromosome 7 gain, chromosome 9p loss and chromosome 10 loss [11,12,1]. The inferred cell components here all show gain of chromosome 7 and loss of chromosome 9p, suggesting these are early events in the tumor’s evolution. Four of the inferred components also show loss of chromosome 10 (Fig. 6, (a)). In addition to the frequent aberrations in chromosome 7, 9 and 10, other chromosomes also display gain (e.g., chromosome 1, 3, 19, 20) and loss (e.g., chromosome 6, 11, 13, 14). There is, however, some notable clonal heterogeneity, similar to inter-tumor heterogeneity observed in systematic studies of GBM [11,24].

**Fig. 6.**
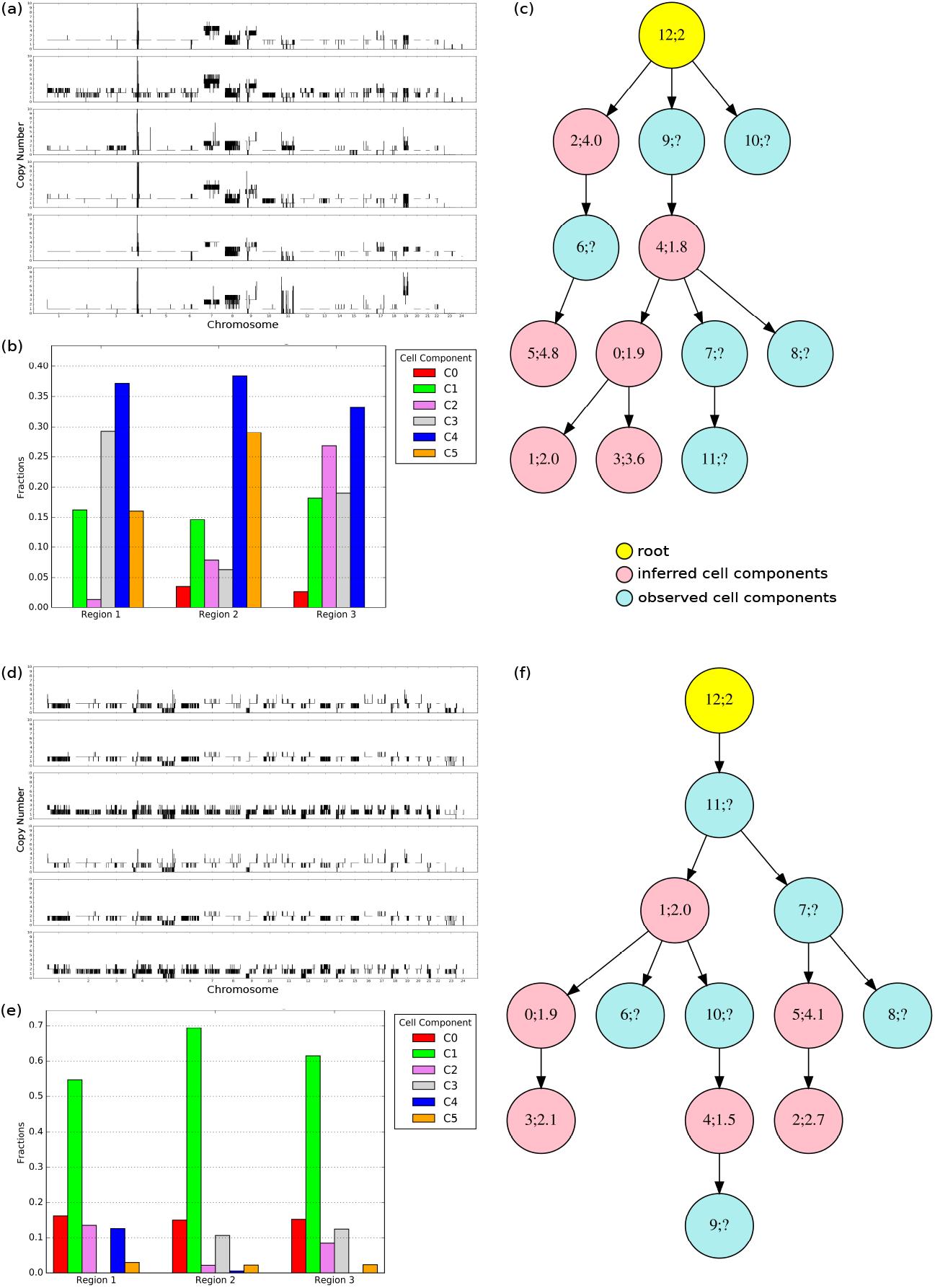
Application on real GBM07 ((a)-(c)) and GBM33 ((d)-(f)) cases. (a)(d) The copy number of each chromosome in inferred cell component C0 (top) to C5 (bottom), axis indicate the chromosome, 1-22 represent autosome, 23 represents X chromosome and 24 represents Y chromosome. (b)(e) The corresponding fraction of each inferred cell component. (c)(f) The phylogenetic relationship among the observed cell components (pink) and inferred cell components (light blue).

Several studies have shown that WGD occurs in about 25% of glioblamstoma cases [7,4,5] and have suggested that it is an early event when it occurs. Our model for the GMB07 tumor supports an inference of two distinct WGD events on distinct cell lineages: an early WGS in the transition from components 12 to 2 and a late WGD event in in the transition from component 0 to 3. This inference that there are multiple WGD events depends on having both sequence data supporting the tree topology and FISH data supporting the specific ploidy changes and therefore supports the value of the miFISH analysis in providing more direct measurements of ploidy and allowing sampling of larger numbers of cells, and thus better detection of rarer clones. Although we do not have information about the ploidy for the observed cell components, we may infer them based on the fact that the components with similar ploidy tend to occur in the same branches on the tree (Fig. 5 (c)). A manual maximum parsimony imputation of WGD events suggests that observed component 6 is most likely tetraploid and all other observed components are most likely diploid.

Prior pan-cancer studies have suggested that WGD often touches off a cascade of more localized CNA losses, with particular marked chromosome losses [37]. Pronounced focal CNA is evident in all of our inferred components. For example, we find that component 1 exhibits widespread CNA events with losses occurring in chromosome 9p,while component 2 and 5 exhibit more obvious losses in chromosome 9p and 10 after WGD (Fig. 6, (a); node 1, 2, 5 in (c)).

Studying mixture fractions of the inferred clones provides additional insight into likely GBM progression. We found that component 4 (blue bar in Fig. 6 (b)) has a relatively large proportion in all three regions, suggesting this might be closer to an ancestral population from which the tumor as a whole arose. That is consistent with the finding that chromosome 7 gain and chromosome 9p loss found in this component are early CNA events in the tumor and perhaps key drivers of tumorigenesis. Noticeable proportions of components 2, 3, and 5 are found in at least in one region (pink, grey and orange bars in Fig. 6 (b)) but with sizable differences by region. This inference is again consistent with the idea the tumor has been shaped by multiple distinct WGD events, with different regions of the tumor dominated by cell lineages tracing to different WGD events.

Fig. 6 (d)-(f) show the results for the GBM33 case. Although we observe some similar events to GBM07, such as (partial) gain in chromosome 7 and loss in chromosome 9p and partial loss in chromosome 10, the global pattern of CNA is quite different. First, GBM33 exhibits a pattern more dominated by focal CNA rather than chromosome-scale changes. Second, GBM33 shows less extreme change at sites of high amplification than does GBM07 even where they amplify common loci (e.g., large copy numbers in chromosome 4, Fig. 6, (a), (c)). Third, there seems to be less tetraploidy or WGD events in GBM33 ((f) in Fig. 6). A single clone 5 is inferred to be approximately tetraploid and it is inferred to be ancestral to a single approximately triploid clone 2, consistent with a recurrent pattern of transition from tetraploid to pseudotriploid observed in past miFISH study of WGD-prone tumors [26]. We can infer that the observed single cells are largely diploid, although it would be ambiguous in a maximum parsimony analysis whether clones 7 and 8 are diploid (corresponding to a single WGD in the descent from 1 to 7) or tetraploid (corresponding to a single WGD in the transition from 7 to 5). Fourth, GBM33 overall shows less pronounced clonal heterogeneity, with the single diploid clone 1 dominant in all three tumor regions. Notably, the tetraploid clone is inferred to be fairly rare, with the pseudo-triploid clone slightly more common but still minor. The quite different reconstruction in the case of GBM33 versus GBM07 indicates that the method is sensitive to variations in profiles of CNA tumor-to-tumor.

## 4 Conclusions and Discussion

We have extended tumor phylogeny methods to incorporate copy number measurements by DNA-FISH, in addition to bulk and single-cell sequence data, as a source of more precise measurements of tumor ploidy and clonal frequencies. The results show that each source of data contributes separately to a more accurate picture of copy number evolution in cancers, with the combination of all three data types yielding improved accuracy in resolution of whole-genome copy number profiles. We demonstrate by application to a pair of glioblastoma cases that the new methods can provide novel insight into the role of copy number evolution in single cancers. The results suggest the value of supplementing sequence data with additional data sources such as miFISH in accurately reconstructing evolution by CNA mechanisms in tumors exhibiting chromosome instability.

The work described here suggests a number of avenues for further research. One limitation of our method is that few tumors currently are studied by the combinations of technologies examined here. We suggest that it will be enlightening to conduct further studies where sequence is paired with miFISH, or perhaps alternative methods providing similar ability to estimate ploidy and/or clonal frequency more accurately, particularly for understanding evolution in cancer types prone to chromosome instability and aneuploidy. Second, the present work, like our prior work [21], suggests the value of an accurate single-cell phylogenetic model in improving deconvolution. Accurately reconstructing evolutionary trees in copy number space, even with known single-cell data, remains a challenging problem. While there is prior theory for reconstructing copy number evolution [9,15], no models are comprehensive for all of the mechanisms of CNA evolution we know about and developing comprehensive models that are scalable to large single-cell, whole genome data remains a challenge. There are also many other alternative technologies that might be incorporated into the mix of multi-omic data to improve phylogeny inference (e.g., long read or linked read sequencing, single-cell RNA-seq and bulk RNA-seq) that have been considered in other work (e.g., [31]) and might provide other synergistic advantages for the present problem. In addition, there is likely room for improvement in better solving the central optimization problem of our work. Additional data (omitted for space) shows that increasing the number of rounds of optimization from 10 to 100 frequently leads to improvement in the objective function, although this improvement translates into negligible change in mean accuracy and RMSD measures. This observation suggests potential for improvement in both the definition of the objective function, to better match true solution quality, and in the algorithms, for efficiently solving for the objective.

## Acknowledgements

This research was supported in part by the Intramural Research Program of the National Institutes of Health, National Library of Medicine and both Center for Cancer Research and Division of Cancer Epidemiology and Genetics within the National Cancer Institute. This research was supported in part by the Exploration Program of the Shenzhen Science and Technology Innovation Committee [JCYJ20170303151334808]. Portions of this work have been funded by U.S. N.I.H. award R21CA216452 and Pennsylvania Dept. of Health award 4100070287. The Pennsylvania Department of Health specifically disclaims responsibility for any analyses, interpretations or conclusions.

## A Supplementary Material

### A.1 Supplementary Methods

#### A.1.1 Coordinate descent for deconvolution

The original deconvolution problem shown in Eq. 1 is non-convex, and it is hard to derive a closed form for the solution, so we apply a coordinate descent method to solve **F, S, C, P** iteratively by following the order of Sec. 2.2 to Sec. 2.5 with the corresponding constraints for each term (Algorithm 1).

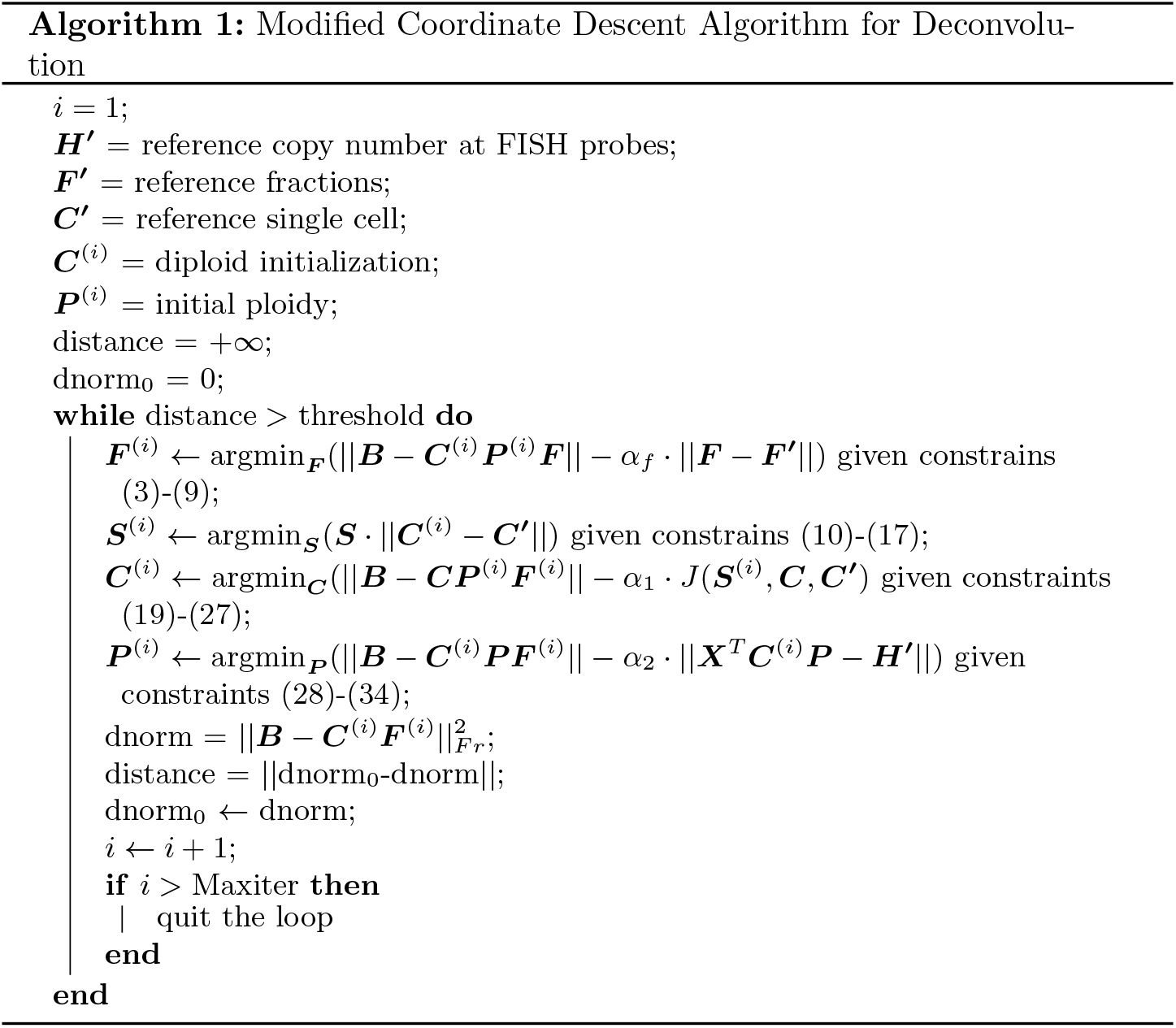

#### A.1.2 Extending the reference FISH matrix

The original *Index* contains the indices of the 8 original FISH probes that in the SCS data. However, compared to the 9934 genomic positions in the SCS data, 8 probes only contribute a very tiny portion to the copy number inference. Alternately, we find that the genomic positions around the FISH probes are highly correlated (Fig. S1), then we extend the *Index* by adding to it the consecutive genomic positions that are highly correlated with the FISH probes (light blocks in the Fig. S1, threshold=0.95).We use two pointers to make sure the correlated genomic positions are consecutive to each other and to the FISH probe (Algorithm 2), and those positions that may be also highly correlated but far away in the genomic positions or even in the different chromosomes would not be considered as correlated.

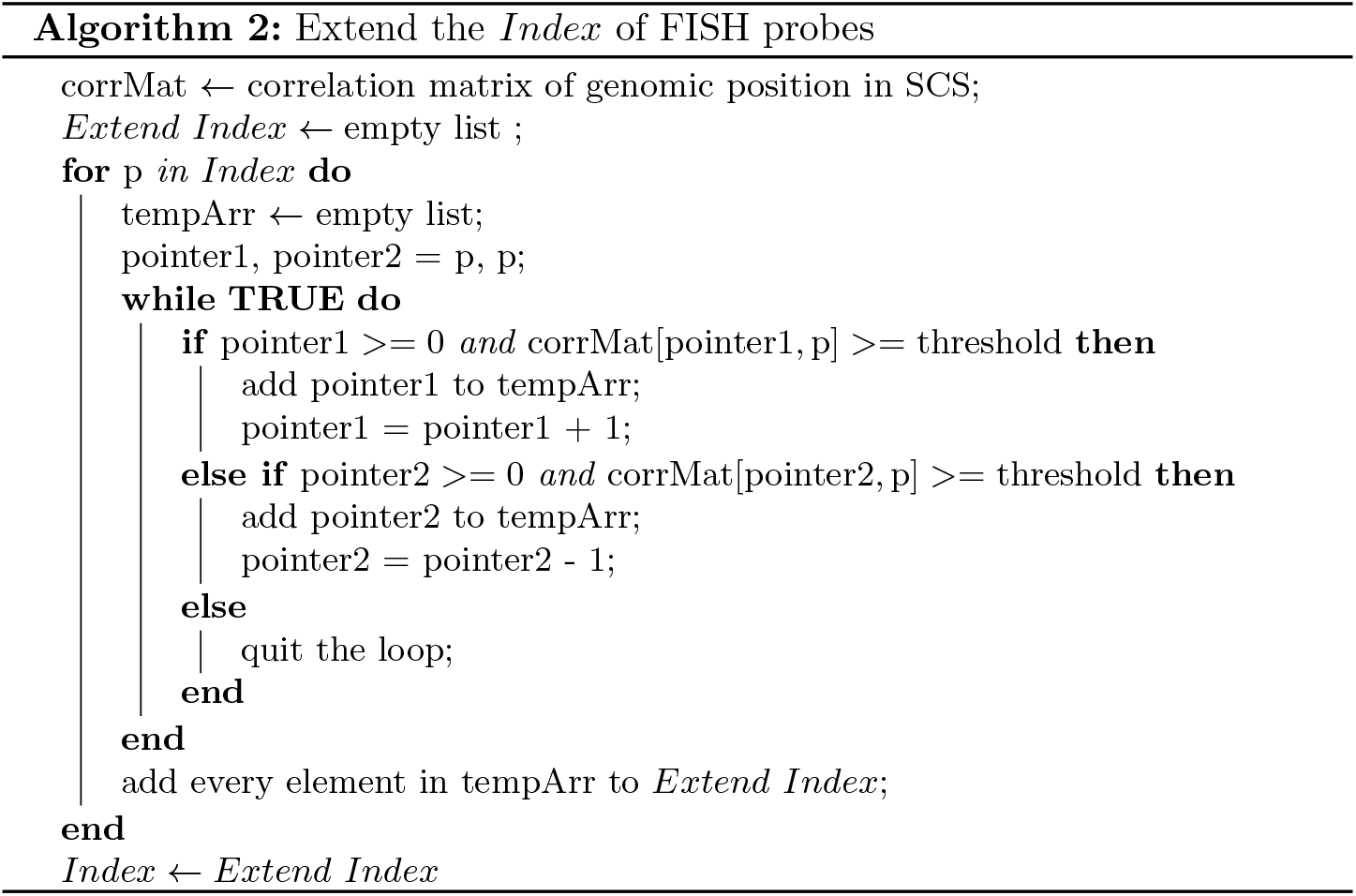

#### A.1.3 Semi-Synthetic Data Simulation

This section describes our protocol for simulating data to test the algorithms. The guiding principle of the method is to generate a ground truth dataset, in which all properties are known and resemble the GBM data, then subsample artificial bulk, SCS, or FISH data from that single ground truth. We set NUM_REGI0NS=3 and NUM_PR0BES=8 and MAX_C0PY=10 to match the GBM data. We define this ground truth in terms of six data structures:

1. 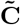: a matrix of normalized copy number profiles of all selected cells, including major, minor and tiny clones. Each column of **Ĉ** corresponds to a ground truth single cell and each row to the mean copy number at a single genomic locus, where it is assumed the rows collectively span the full genome. We assume each cell (column) is normalized to mean diploid count.
2. **Ĉ:** a matrix of normalized copy number profiles of major clones in each tumor region. According to previous description, **Ĉ** was generated by picking the first two components in 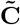 and used to calculate copy number accuracy and RMSD for performance estimation.
3. 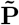: a diagonal matrix of half ploidies, where each non-zero element 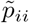 provides a scaling factor to convert the diploid row 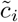 to absolute (unnormalized) copy numbers.
4. 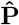: a diagonal matrix of half ploidies, where each non-zero element 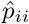 provides a scaling factor to convert the diploid row *ĉ_i_* to absolute (unnormalized) copy numbers.
5. 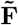: a matrix of mixture fractions, where each row corresponds to a selected cell and column defines a probability density describing frequency of occurrence of each cell type in the bulk samples.
6. 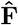: a matrix of mixture fractions, where each row corresponds to a major clone and column defines a probability density describing approximate frequency of occurrence of each major clone in the bulk samples. 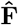 is derived from 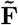, but column is also normalized to 1.

**Fig. S1.**
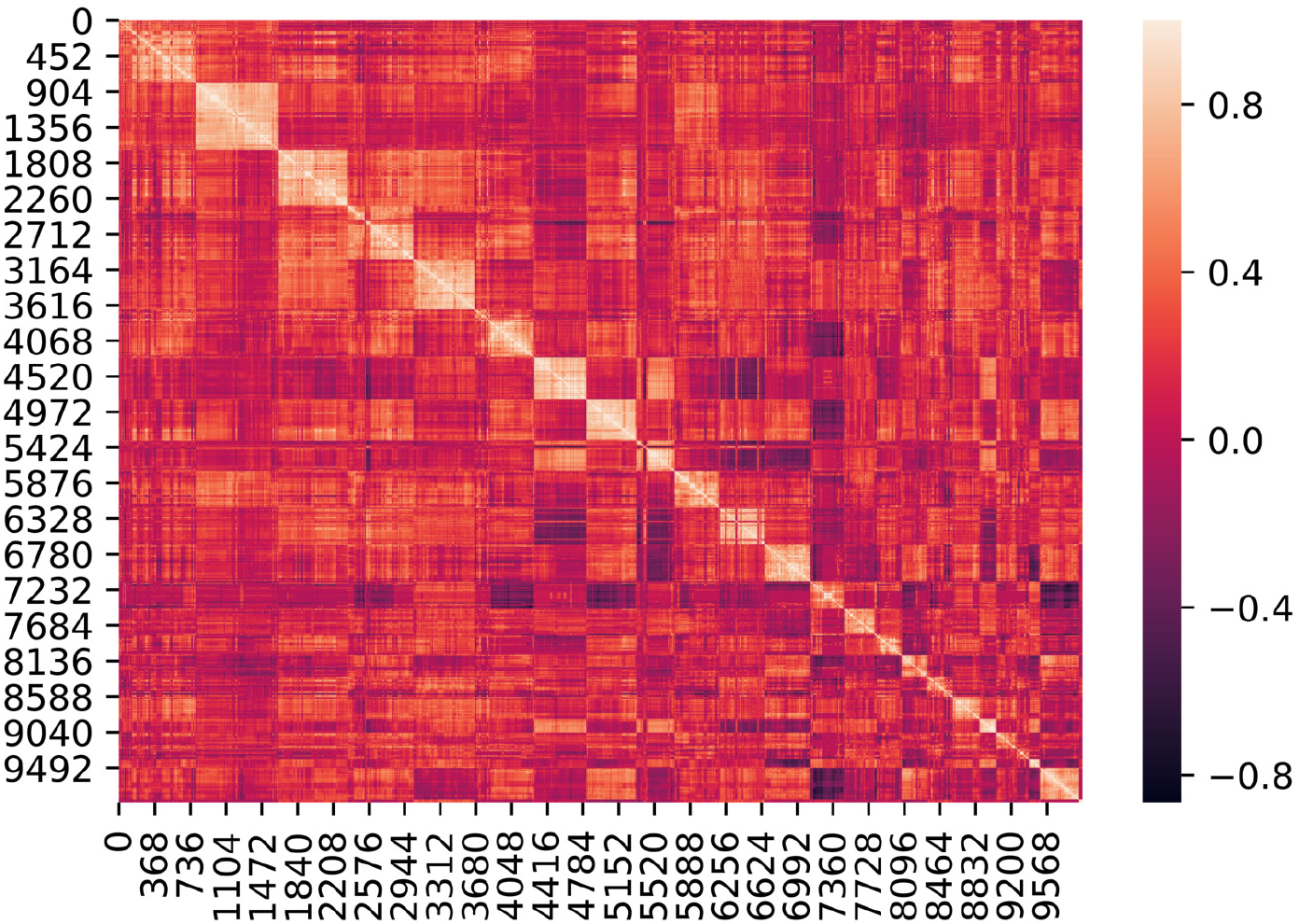
Correlation matrix for genomic positions. The light blocks indicates the neighbouring genomic position are highly corrleated in positive direction. For each one of 8 original FISH probe indexes, We search the consecutive genomic positions that are highly correlated with it and add it to *Index,* so that we extend the original *Index* from length of 8 to the length around 100 (please also refer to Fig. S2, step (9)).

We first define this ground truth model, then generate simulated data of each needed type by sampling from the model. These processes are described step-by-step below.

**Fig. S2.**
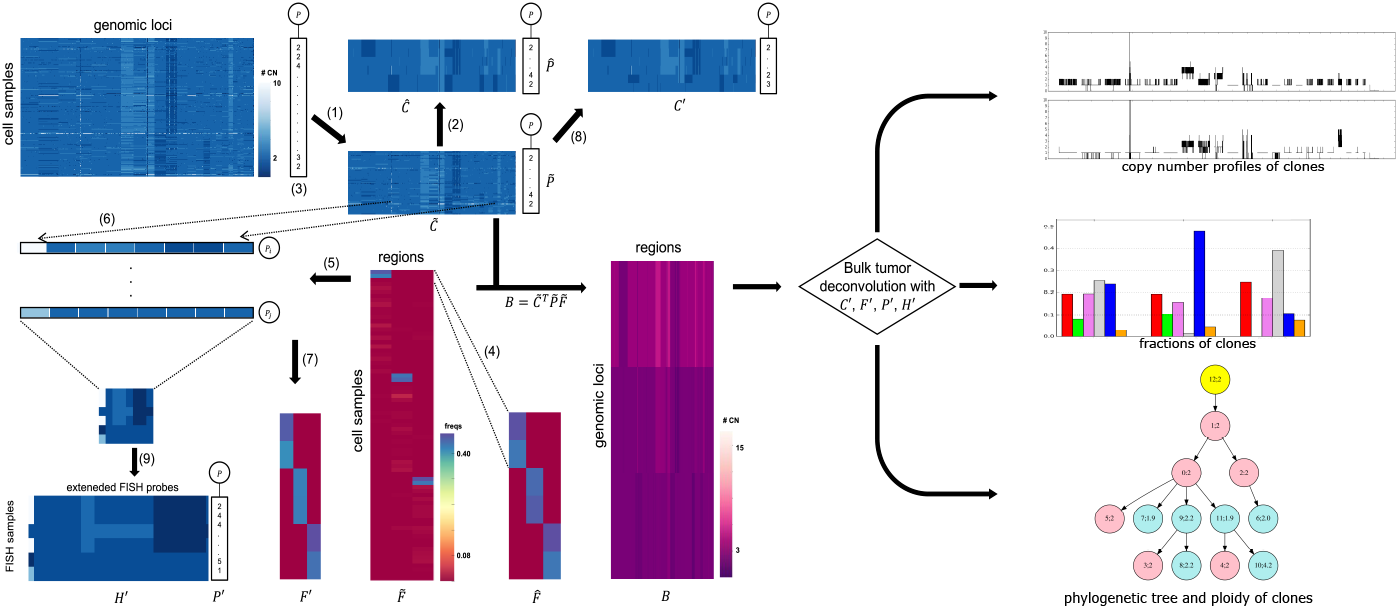
Workflow of the simulation and deconvolution. The figure shows the process from real SCS data to select SCS clones, sample ploidies, simulate mixture fractions, simulate FISH and simulate bulk genomic data. We then deconvolve the bulk data into copy number profiles of a set of inferred clones each with a defined ploidy and set of mixture fractions across tumor regions, as well as a phylogenetic tree relating these clones. We then compare these outputs with the ground truth data to evaluate our model. Further methodological details are provided in the text. Note that the images in this figure are purely illustrative and do not show true data from any particular analysis.

*Selecting clones from SCS data* We first select copy number vectors to instantiate the normalized copy numbers in 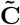 and identify these as clones of the model. We use the true SCS data for this purpose. We uniformly at random select 25 single cells from each of the NUM_REGIONS regions to have 75 cells in total, of which the copy number and ploidy make nonzero contribution in the simulated bulk tumor sample later. The true copy number data of the selected cells define the columns of 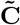. Of the 25 single cells from each region, we denote the first 2 as *major* clones or high-frequency clones, and the remaining 23 cells as *minor*, or low-frequency, clones for that region. For each region, we model the assumption that, within the tumor, cells from the other occur but with very small frequency. Thus for each region, we designate the 50 cells from the other two regions as *tiny* clones, which will let these cells effectively serve as noise in the analysis (Fig. S2, step (1)). The two major clones from each regions to compose **Ĉ**, which has 6 clones in total (Fig. S2, step (2)).

*Sampling ploidies* Since the real single-cell sequencing data have been normalized, the ploidy profiles for all samples have been set to 2 (diploidy) by default, and we call them *normalized cell.* The *normalized cells* are a standard target to study tumor evolution, however, the ploidy information is also important during tumor evolution [13,4]. Since we do not know the correspondence between ploidies and WGS copy number vectors in the ground-truth data, we simulate a ploidy for each cell. We sample a ploidy independently for each ground truth cell. We give each ground truth cell *i* a probability *β*_1_ of being diploid, corresponding to 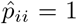. We give it a probability *β*_2_ of tetraploidy, corresponding to 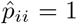. We then allow a probability *β*_3_(= 1 — *β*_1_ — *β*_2_) of some other ploidy, selected uniformly from [1, 3, 5, 6, 7, 8]. Currently, *β*_1_ = 60%, *β*_2_ = 30%, *β*_3_ = 10%. Thus, at present:

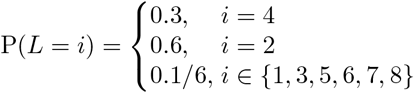

where *L* represents the ploidy number for and *P*(*L*) is the probability of each ploidy number, then we have an additional tag of ploidy number for each SCS sample (Fig. S2, step (3)).

*Simulating mixture fractions* We next assign mixture fractions 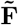 to the clones. We follow our previous work [21] to use a Dirichlet distribution *Dir*(*γ*), to assign multinomial frequencies to clones selected as in A.1.3. *γ* is a vector of concentration parameters that allows different cell components to have different contributions in the bulk tumor. The vector *γ* is generated to model that in the Dirichlet distribution, all regions have a equal prior probability of contributing to the bulk tumor. Following our previous work [21], for each region, we set *γ* to be 100 for these major clones, 1 for the these minor clones and 0.01 for these tiny clones. Because there are three regions, we take the sum of the three vectors *γ*, one for each region, and use the sum as the parameters to the Dirichlet distribution. Then we retrieve the simulated fractions of major clones to compose 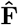, and normalize each column to 1 (Fig. S2, step (4)), this would be used as fractions RMSD calculation later.

*Simulating bulk genomic data* Once we have defined a ground truth dataset, we simulate each source of input data for a given problem instance from this common ground truth. We first simulate bulk data from the reference model by assuming that each regions samples all clones from their ground truth proportions and with the ground truth copy number vectors and mixture fractions. That is, we simulate the input bulk matrix ***B*** as 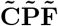.

*Simulating FISH copy number profiles* We next simulate FISH data using the genomic positions of the same NUM_PROBES loci as in the real data. Because we require the ground truth mapping of simulated FISH to whole-genome copy number vectors, we do not use true FISH probe counts or assigned ploidies for this simulation. We assume known absolute genomic positions of *S_begin_* and *S_end_* of each genomic interval in 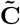 and absolute genomic loci *H_begin_* and *H_end_* of all FISH probes according to the reference genome hg19. This provides us a way to retrieve corresponding copy number as the copy number of probes in FISH. We also save an array *Index* mapping overlaps of SCS intervals and FISH probes for later use.

To simulate a FISH cell in a region, we use the two major clones for the region and restrict their copy numbers to the intervals overlapping the FISH probe. If the interval for a given FISH probe is included in a given SCS interval, we assign the FISH probe count to be the copy number of the corresponding SCS interval. If the FISH probe crosses two SCS intervals, we assign the FISH probe count to be a weighted average of the copy numbers of the two SCS intervals, weighted by the length of the FISH probe in each SCS interval (Fig. S2, step (6)). No FISH probe covers more than two SCS intervals in the real data, so we do not consider any other cases.

We also optionally randomly perturb copy numbers to simulate errors in FISH probe counts before transferring them to be unnormalized. This can be represented in terms of noise parameter *q_f_*, where with probability *q_f_* a probe count will be increased by 1, with probability *q_f_* it will be decreased by 1 unless already zero, and with probability 1 — 2*q_f_* it will be unaltered. Both before and after adding noise, the FISH copy numbers are capped at MAX_COPY.

We repeat this process for 1000 FISH cells in each of NUM_REGIONS tumor regions to generate a simulated FISH data set (Fig. S2, step (5)).

Simulating FISH frequencies We assume that the FISH data provide an approximate measure of the distribution of clonal fractions. From the 1000 FISH cells simulated in A.1.3, we calculate the fraction of each FISH cell for each region by calculating the proportion of each FISH cell out of the total number of FISH cells (1000), and then extract the fractions of the first two largest clones from each region. We combine these fractions and allow the sum of each column to be less than 1, since in real data, it is possible that there would be a small proportion of cells that are not represented by the major clones. Then the resulting fraction matrix **F′** represents the fraction of each major clone across the FISH cells for each region, which can be used as reference for the fractions of the major clones in SCS data (Fig. S2, step (7)).

Simulating SCS data To simulate a set of SCS data, we select cells independently at random from 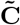 with probabilities for each cell in each region as defined in 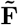. The resulting SCS matrix **C′** would then consist of normalized single cells, where each column of **C′** initially corresponds to some column of **Ĉ**, allowing for repetition (Fig. S2, step (8)).

We further allow the data to be perturbed by a noise model with parameter *q_s_*, where with probability *q_s_* each copy number will be increased by 1, with probability *q_s_* it will be decreased by 1 unless already zero, and with probability 1 – 2*q_s_* it will be unaltered. Also, we would not allow for copy number to exceed 10 after perturbing the noise.

### A.2 Supplementary Results

#### A.2.1 Deconvolution without SCS Data

We initially tested our model in the scenario where we do not have real SCS data but only have FISH available. To incorporate the tree part of the objective function, we make an artificial reference cell matrix with all diploid copy number for every entry. We do the same process as described in Sec. 3. Fig. S3 shows the average result. From the top to bottom are the results without noise and without ploidy change, with 10% but without ploidy change, without noise but with ploidy change, with 10% and with ploidy change, respectively. We can find that, compared to the results in Fig. 1 and 3, the performance was worse for most of the cases. This observation suggests that the real SCS data plays an important role in the reference, which is consistent with the conclusion of our previous work [21]. However, this loss is not obvious if we do not perturb the ploidy, as assuming diploid reference cells effectively provides an informative prior probability for the inference. When we implemented the change of ploidy, the difference of performance with and without real SCS become evident. Nonetheless, the addition of FISH data still substantially improves accuracy relative to inference from bulk sequence data alone.

#### A.2.2 Deconvolution with different number of iterations

As mentioned in 3.1.1, our current model reduced the maximum number of iterations for the Gurobi solver from 100 to 10 relative to our earlier work, as we found that increasing the number of iterations could greatly increase run time while generally not significantly improving our quantitative measures of performance. Here, we evaluate the effects of this change by showing performances with two different maximum numbers of iterations (Fig. S4) in the case of 10% noise and with variable ploidy 3.1.2. In all cases, optimization can terminate before the maximum number of iterations based on the convergence test of Algorithm 1. The cyan box shows the results of maximum iteration = 10 and violet box shows the results of maximum iteration = 100. We can see that though there is some variation between each pair of results, the average values show no consistent pattern of improvement with increasing numbers of iterations and no significant difference between the two. While additional rounds of optimization do sometimes lead to better solutions, the results suggest that improvement is generally small and that further refinement of the objective function does not reliably translate to better solutions as assessed by our performance measures.

**Fig. S3.**
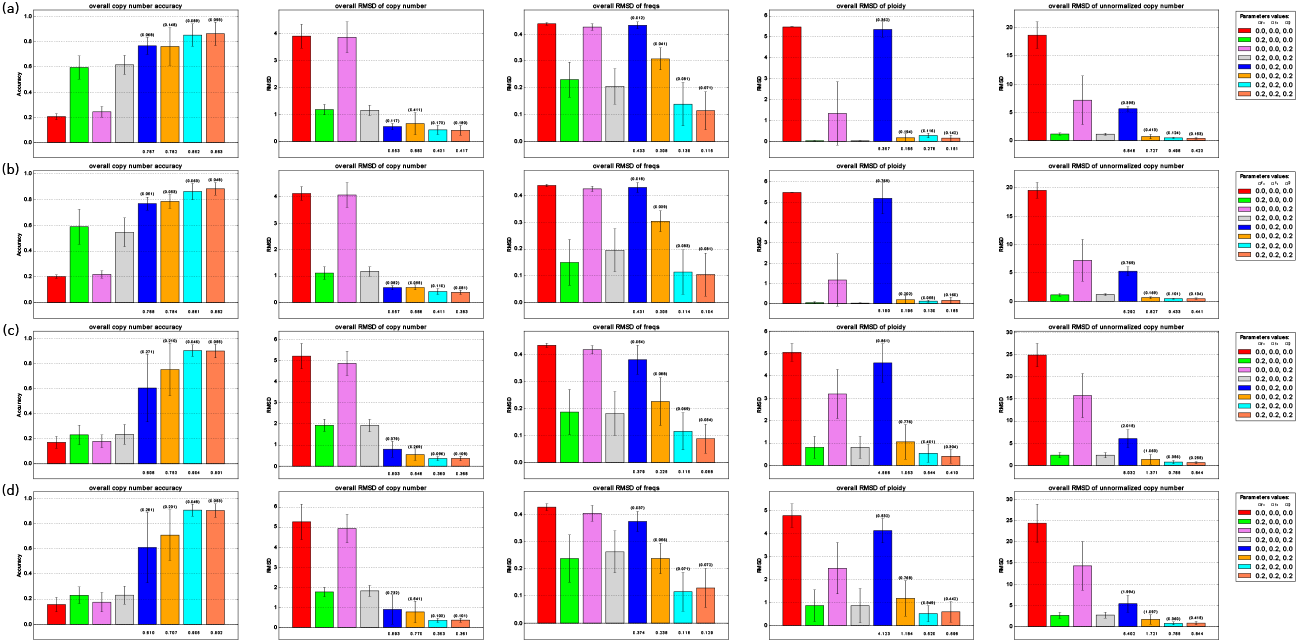
Average accuracy and RMSD of the deconvolution without real SCS data. From (a) to (d) are the results without noise and without ploidy change, with 10% noise but without ploidy change, without noise but with ploidy change, and with 10%) and with ploidy change, respectively. In each subplot (a) to (d), the barplot shows the average accuracy of copy number, average RMSD of copy number, average RMSD of fraction and average RMSD of ploidy. All the labels are the same and the numbers represent the value of *α_f_*, *α*_1_, *α*_2_, which are regularization terms for ||***F — F′***||, *J*(***S,C,C′***) and ||***X^T^CP – H′***||, respectively. 0.0 means the corresponding term is not included in the model

**Fig. S4.**
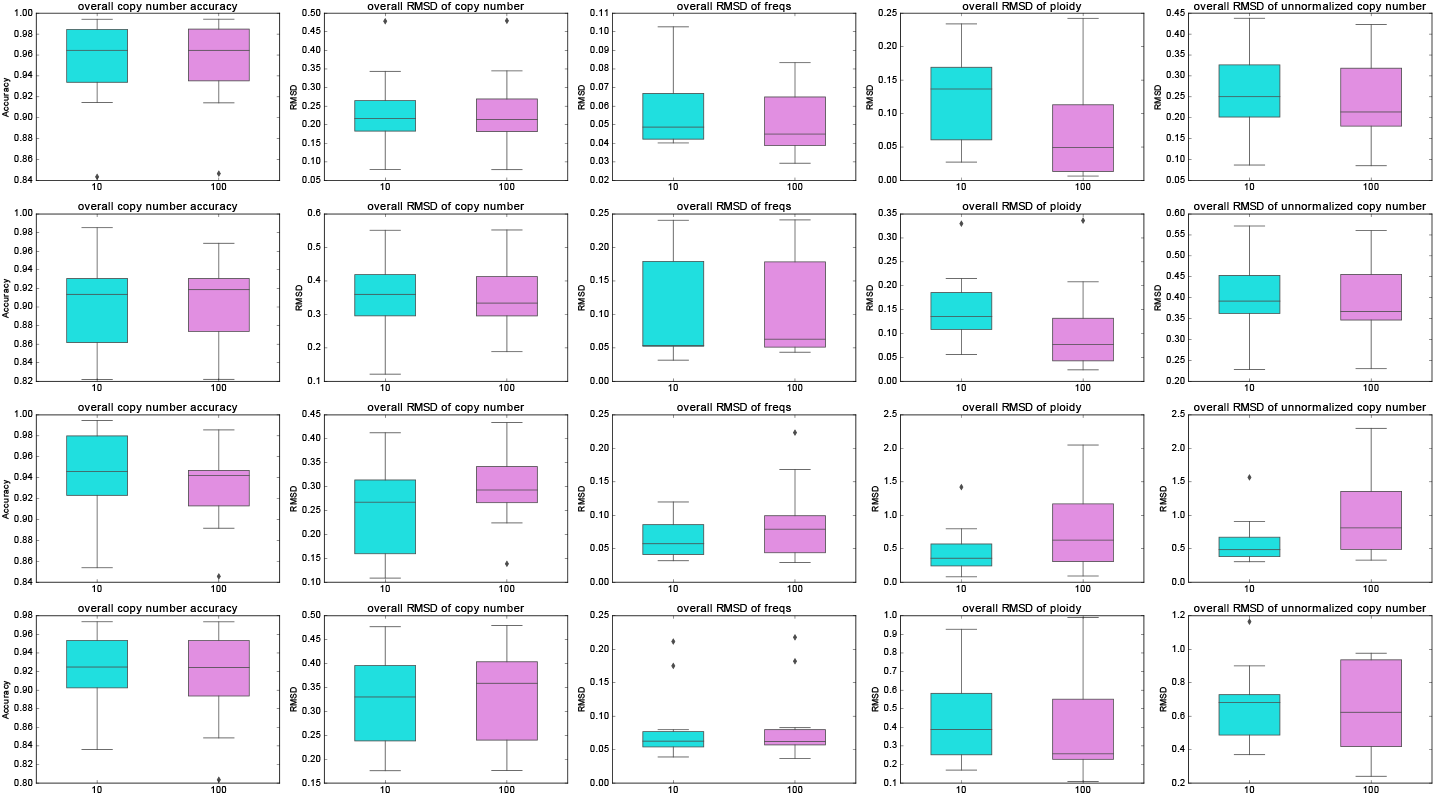
Performance comparison for varying numbers of maximum iterations of optimization. This figure shows differences in performance for 10 versus 100 maximum iterations of optimization in the case of 10% noise and variable ploidy. Cyan boxes represent results of maximum iterations = 10 and violet boxes represent results of maximum iterations = 100. From the left to the right, we present the performance comparison in overall copy number accuracy, overall RMSD of copy number, overall RMSD of frequency, overall RMSD of ploidy and overall RMSD of unnormalized copy number, respectively.

#### A.2.3 Sensitivity to parameters change

In the previous sections, we turned on or off the three parameters (*α_f_*, *α*_1_, *α*_2_) by setting them either to 0.2 or 0.0. We chose 0.2 heuristically as a good default value for similar regularizations in our previous work [21]. In this section, we explored the question of sensitivity of the parameters to determine whether the results would be highly dependent on parameter choices. To evaluate this, we performed a parameter scan around the value of 0.2 to test different combinations of the three parameters, focusing specifically on the case of 10% noise and with variable ploidy. In the set of parameter combinations, we found that the model is minimally sensitive to changes of parameters in the measurement of normalized copy number but somewhat more sensitive to the change of parameters in the measurement of frequency, ploidy and unnormalized copy number (Fig. S5). For example, when we fix *α*_2_ to be 0.1, the average performances in each heatmap does not change much in copy number inference (1^*st*^ and 2^*nd*^ columns in Fig. S5) but shows a little bit more oscillation in the rest of measurements when we increase *α_f_* and\or *α*_1_ (3^*rd*^, 4^*th*^ and 5^*th*^ columns in Fig. S5). When we fixed *α_f_* and *α*_1_, we observed that ploidy inference does not reveal a simple pattern of better or worse average performance across different combinations of parameters, which indicates that the parameters may influence performance in a more complicated way. Further, there is no one ideal parameter set for all measures, but rather improvement by different measures with different parameter variations. Nevertheless, the default setting of parameters (*α_f_* = 0.2, *α*_1_ = 0.2, *α*_2_ = 0.2) seems to yield a good consensus that provides a reasonable set of trade-offs in the performance across all the measurements (3^*rd*^ and 4^*th*^ rows in Fig. S5).

**Fig. S5.**
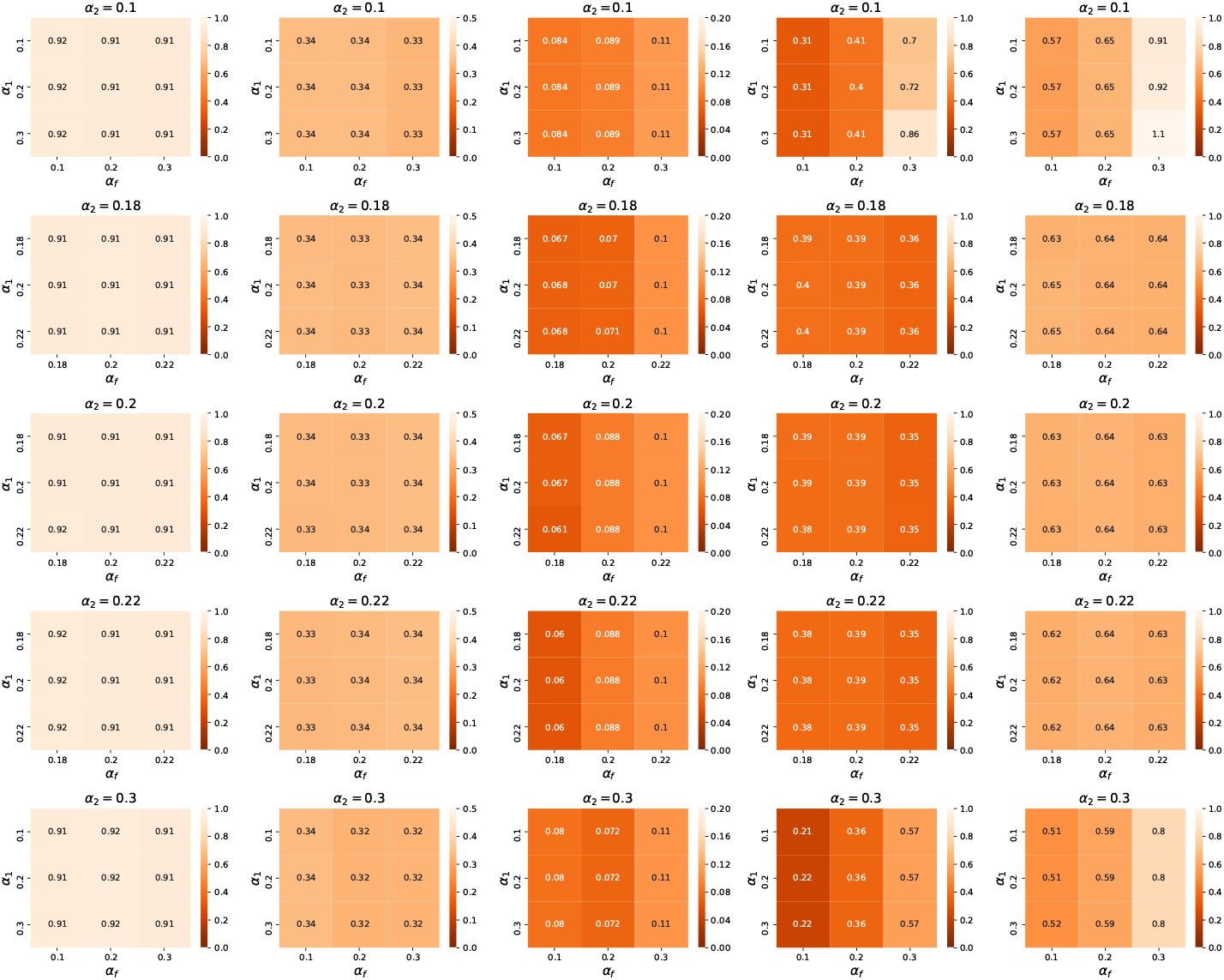
Model performance for different combinations of parameters. This figure shows the results of sensitivity tests on semi-simulated data, where we vary the value of *α*_2_ to be 0.1, 0.18, 0.2, 0.22, 0.3. For each *α*_2_ across subfigures, we provide complete combinations of *α_f_* and *α*_1_ in (0.1, 0.2, 0.3) or (0.18, 0.2, 0.22) in each figure. From the left to the right, we present the performance in overall copy number accuracy, overall RMSD of copy number, overall RMSD of frequency, overall RMSD of ploidy and overall RMSD of unnormalized copy number, respectively. In each case, we model 10% noise and variable ploidy. The value in each block represents the average performance of *n* = 10 experiments.

